# TPM4 condensates glycolytic enzymes to fuel actin reorganization under hyperosmotic stress

**DOI:** 10.1101/2024.07.09.602822

**Authors:** Wenzhong Yang, Yuan Wang, Geyao Liu, Yan Wang, Congying Wu

## Abstract

Actin homeostasis is fundamental for cell structure and consumes a large portion of cellular ATP. It has been documented in the literature that certain glycolytic enzymes can interact with actin, indicating an intricate interplay between the cytoskeleton and cellular metabolism. Here we report that hyperosmotic stress triggers actin severing and subsequent phase separation of the actin-binding protein TPM4. TPM4 condensates glycolytic enzymes such as HK2, PFKM, and PKM2, and adhere to and wrap around actin filaments. Notably, the condensates of TPM4 and glycolytic enzymes are enriched of NADH and ATP, suggestive of their functional importance in cell metabolism. At cellular level, actin filaments assembly is enhanced upon hyperosmotic stress and TPM4 condensation, while depletion of TPM4 impaired osmolarity-induced actin reorganization. At tissue level, co-localized condensates of TPM4 and glycolytic enzymes are observed in renal tissues subjected to hyperosmotic stress. Together, our findings suggest that stress-induced actin perturbation may act on TPM4 to organize glycolytic hubs that tether energy production to cytoskeletal reorganization.

**Graphical Abstract:** 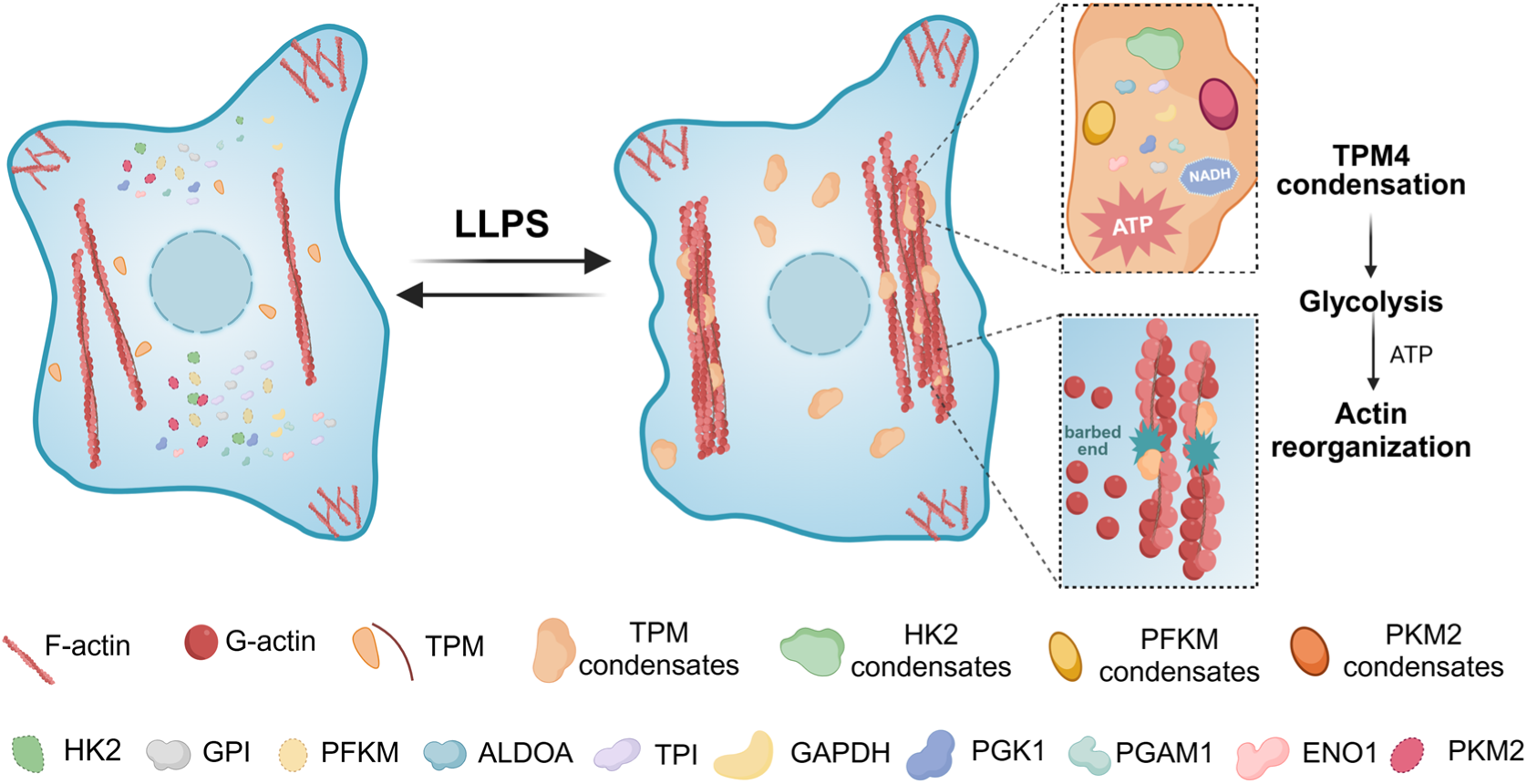

## Introduction

Actin, the most abundant protein in eukaryotic cells, polymerizes through nucleation and elongation from globular actin monomers (G-actin) to form filamentous actin (F-actin). The dynamic rearrangement of the actin cytoskeleton is a highly energy-consuming process. Early studies found that the turnover of F-actin accounted for 50 % of total ATP consumption in unstimulated platelets and neurons[1, 2]. The direct interactions between glycolysis enzymes and actin indicate that glycolysis may provide the necessary fuel for actin rearrangement[3] [4] [5]. Actin cytoskeleton has recently been shown to adjust glycolysis level. Soft substrate induced stress fiber disassembly, releasing E3 ubiquitin ligase tripartite motif (TRIM)-containing protein 21 (TRIM21) to degrade phosphofructokinase (PFK) and thus downregulate glycolysis[6]. Insulin stimulation triggers F-actin remodeling in epithelial cells through the PI3K-PIP3-Rac1 pathway, increasing the amount of actin-free aldolase (ALDOA) in the cytoplasm, enhancing glycolytic flux[7]. Purified PFK protein has long been documented to bind polymerized actin and formed paracrystal in vitro [3]. In yeast, F-actin acts as a scaffold for glycolytic enzymes to form a multi-enzymatic glycolytic complex[8]. It has been hypothesized that actin cytoskeleton can compartmentalize glycolytic enzymes[9]. Functions of the interplays between F-actin and these enzymes including direct interactions and indirect regulation through complex signaling pathways and protein modifications have just started to be elucidated.

Various cellular stress conditions, including hyperosmotic stress, impact cell morphology and metabolism. In turn, cells rewire their cytoskeleton and energy production to survive harsh conditions. During starvation, cells adapt by inhibiting growth, inducing autophagy, and reprogramming metabolism[10] to optimize limited energy supplies, where actin cytoskeleton reorganizes to assist with autophagosome formation[11] and cargo trafficking[12] as well as cell death[13]. Changes in extracellular osmolarity not only influence the life cycle of single cells, but also challenge multicellular organisms by perturbing the cell cortex and trafficking systems, cytoplasm viscosity, nuclear volume and protein stability[14–16]. Notably, hyperosmotic stress induces the condensation of multivalent proteins[17], potentially influencing cellular adaptation. Whether and how phase separation is involved in actin cytoskeletal remodeling under hyperosmotic stress remains unclear.

Protein liquid-liquid phase separation (LLPS) serves as a reaction crucible for biochemical reactions and recruits specific interacting factors, thereby promoting reaction progress[18, 19]. Multienzyme metabolic assemblies have been identified in various tissues and species. For instance, “glucosome” is a reversibly organized assembly of enzymes involved in both glycolysis and gluconeogenesis, regulating glucose flux at the subcellular level in human cancer cells. Glucosomes contain enzymes such as PFK, fructose-1,6-bisphosphatase (FBPase), phosphoenolpyruvate carboxykinase (PEPCK), and pyruvate kinase (PKM). Small-sized glucosomes primarily promote glycolysis, while medium and large-sized ones preferentially partition glucose flux into the pentose phosphate pathway (PPP) and serine biosynthesis respectively[20]. During G1 phase, increased small and medium glucosomes in Hs578T cells suggest heightened glycolysis and PPP, while decreased small glucosomes in G2/M-arrested cells indicate reduced glycolysis in G2 phase[21, 22]. Moreover, under hypoxic stress, yeast and C. elegans neurons form “G bodies” or “glycolytic granules” containing PFK and other glycolytic enzymes. These structures, similar to stress granules and P bodies formed through phase separation[23–28]. The formation of these structures may enhance the activity of the glycolytic pathway, optimize reaction rates, and contribute to ATP production, forming a functional unit known as a “metabolon”[23–25, 29]. Mutants unable to form these structures exhibit decreased cellular viability or impaired synaptic function[24, 25]. Nonetheless, a systematic mechanism for actin cytoskeleton, phase separation and glycolysis remained to be established.

Here we discovered actin-binding protein Tropomyosin 4 (TPM4) can undergo phase separation, forming condensates that adhere to and wrap around actin filaments. These TPM4 condensates play a functional role in recruiting crucial glycolytic enzymes such as HK2, PFKM, and PKM2, promoting glycolysis. Our findings suggest that TPM4 condensates act as a central hub for osmolarity-induced glycolysis upregulation, providing a novel model that links actin cytoskeleton and glycolysis.

## Results

### Dense regions of actin-binding protein contain glycolytic enzymes upon hyperosmotic stress

To visualize actin cytoskeleton in response to hyperosmotic stress, we expressed Lifeact-EGFP to label actin filaments while adding sorbitol to the cell culture medium for the increase of osmolarity. Live-cell imaging revealed that cell body shrank, followed by dynamic actin network reorganization and thickening of individual stress fibers, indicative of hierarchical and coordinated biochemical and biomechanical cellular response to hyperosmotic stress (Figure 1A). Actin assembles actively at the barbed ends. We adapted a barbed end assay[30] to probe de-novo actin polymerization sites. To localize free barbed ends, cells were permeabilized in detergent containing rhodamine-actin at concentrations sufficient to support only barbed end polymerization[31]. We observed that cells exhibited increased barbed ends under hyperosmotic stress (Figure 1B), indicating that hyperosmotic stress may induce actin filament breakage and generation of free barbed ends.

**Figure 1.**
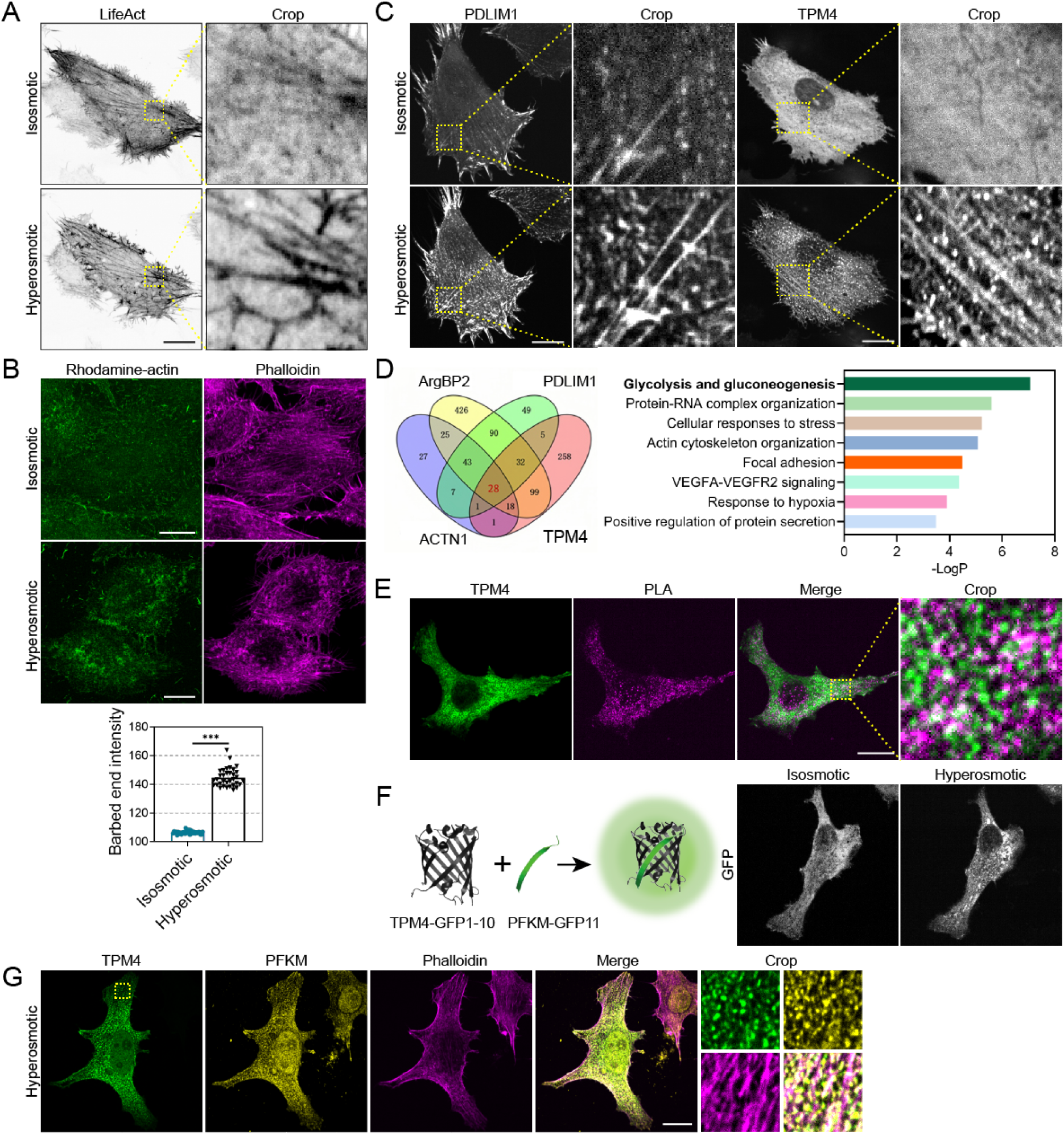
Dense regions of actin-binding protein contain glycolytic enzymes upon hyperosmotic stress. (A) Representative images of MDA-MB-231 cells expressing LifeAct-EGFP before (isosmotic) and after (hyperosmotic) 100 mM sorbitol treatment for 1 h. Scale bar, 20 μm. (B) Barbed end assay showing the distribution of rhodamine-actin before (isosmotic) and after (hyperosmotic) 100 mM sorbitol treatment for 3 min in MDA-MB-231 cells. Actin filaments are labelled by phalloidin. Scale bar, 20 μm. (C) Representative images of MDA-MB-231 cells expressing PDLIM1-AcGFP or TPM4-AcGFP before (isosmotic) and after (hyperosmotic) 100 mM sorbitol treatment for 3 min. Scale bar, 20 μm. (D) Left: venn diagram illustrating the identified proteins by TurboID proximity labeling/Mass-spectrometry using ArgBP2, PDLIM1, ACTN1 and TPM4 fused with TurboID. Right: Enrichment analysis by Metascape showing top enriched signaling pathways. (E) PLA assay showing interaction between TPM4-AcGFP and PFKM under hyperosmotic condition (100mM sorbitol for 3 min) in MDA-MB-231 cells. Scale bar, 20 μm. (F) Left: schematic diagram of split-GFP system expressing TPM4-GFP 1-10 and PFKM-GFP 11. Right: representative images of MDA-MB-231 cells expressing split-GFP constructs before (isosmotic) and after (hyperosmotic) 100 mM sorbitol treatment for 3 min. Scale bar, 20 μm. (G) Representative immunofluorescence images stained with PFKM antibody (yellow) and phalloidin (magenta) in MDA-MB-231 cells expressing TPM4-AcGFP under hyperosmotic condition (100 mM sorbitol for 3 min). Scale bar, 20 μm.

These observation leads to the intriguing question: how do cells regulate actin reorganization under hyperosmotic stress? Actin-binding proteins are known to regulate the actin dynamics, organization, and stability of actin filaments by controlling monomeric availability, nucleation, elongation, and depolymerization[32]. Interestingly, we observed increased localization various actin-binding proteins such as TPM4 and PDLIM1 on actin filaments in response to hyperosmotic stress (Figure 1C). These findings suggest that the remodeling process of actin filaments under hyperosmotic stress may involve the recruitment of multiple components. To investigate the recruited components and their functions in this process, we performed TurboID proximity labeling[33]/Mass-spectrometry using four major actin-binding proteins (ACTN1, PDLIM1, ArgBP2, and TPM4), which have been reported to form dense bodies[34–36] on actin filaments or function as side-binding proteins along actin filaments. Analyzing the results of the four identified groups of mass spectrometry, we found 28 proteins shared by all four groups. Subsequent enrichment analysis using the “Metascape” platform revealed that the “glycolysis and gluconeogenesis” pathway ranked highest in significance (Figure 1D), and among them resides the crucial rate-limiting enzyme PFKM (Phosphofructokinase muscle type) involved in glycolysis.

Actin polymerization demands for high energy consumption. Glycolysis is a quick ATP-producing pathway. It has been documented that purified PFK can interact with actin filaments in vitro[3]. However, the role of actin filaments in regulating PFK in the complex cellular environment remains unclear. We then wondered whether actin binding proteins such as TPM4 may moonlight to facilitate energy support for reorganizing actin cytoskeleton under hyperosmotic stress. To answer this, we first investigated whether TPM4 could interact with glycolytic enzymes. Through the proximity ligation assay (PLA) technique, we identified positive signals resulting from the interaction between TPM4 and PFKM under hyperosmotic stress (Figure 1E). In addition, we employed split-GFP system, wherein TPM4 and PFKM were respectively fused to distinct fragments of the fluorescein moiety of GFP. When shifting cells to hyperosmotic medium, we instantaneously detected enhanced formation of green fluorescent foci (Figure 1F). Furthermore, PFKM showed strong co-localization with TPM4 under hyperosmotic stress, and these colocalized sites are closely associated with actin filaments (Figure 1G).

Together, we detected that hyperosmotic stress promoted an upsurge in actin filaments assembly, and the actin-binding protein TPM4 showed enhanced localization along actin filaments with glycolytic enzymes such as PFKM.

### TPM4 can form condensates through phase separation

When we expressed TPM4-AcGFP in cells, we observed that TPM4 in different morphology - lining with the actin filaments, diffusive, or condensated in the cytoplasm (Figure 2A left panels), the last of which hinted a propensity to undergo phase separation or transition. Prediction analysis revealed that ∼60% region of TPM4 were intrinsically disordered regions (IDRs) (Figure 2A right panels), indicating its intrinsic capacity of phase separation.

**Figure 2.**
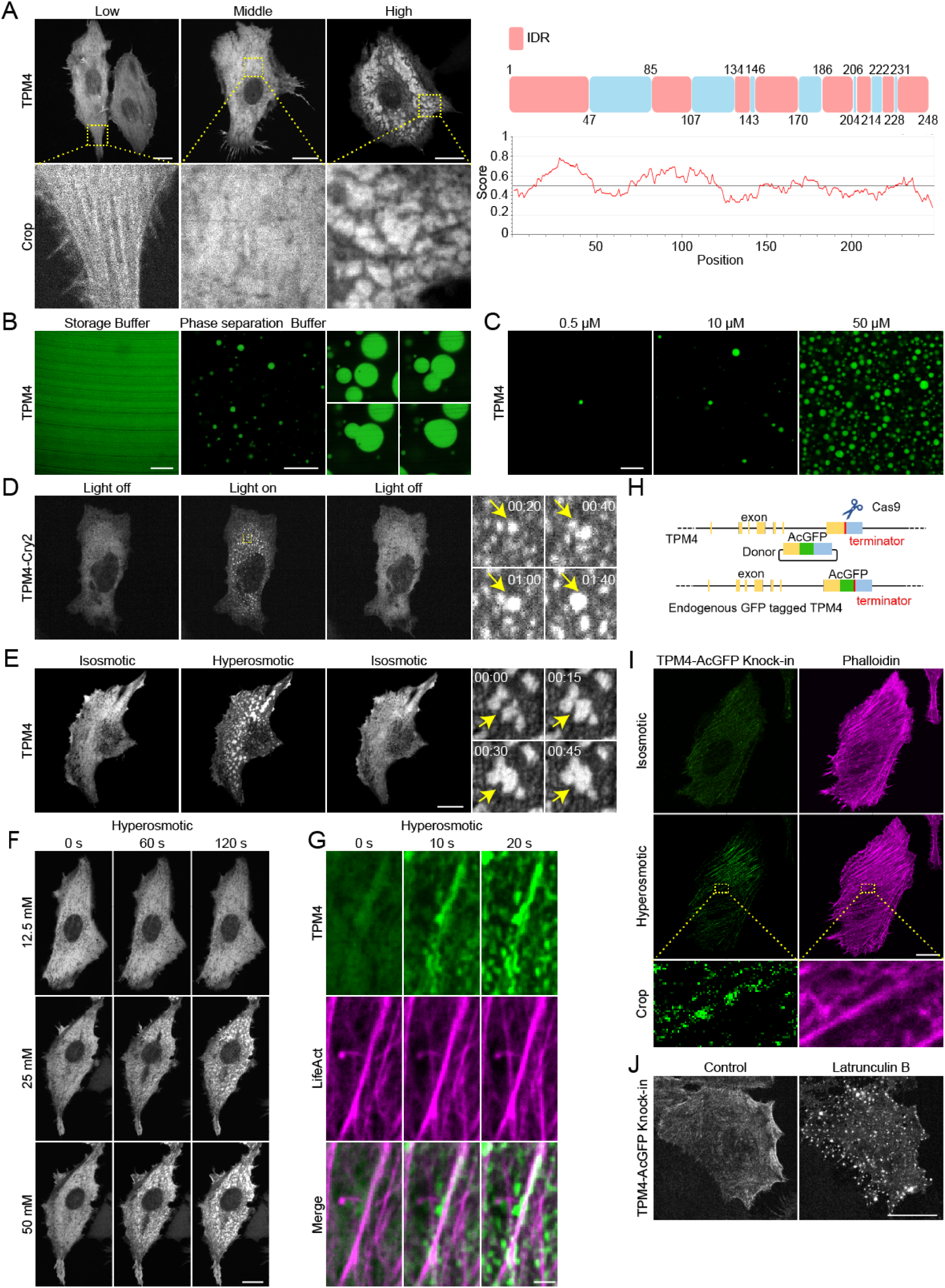
TPM4 can undergo phase separation. (A) Left: representative images of MDA-MB-231 cells expressing low, medium and high level of TPM4-AcGFP. Scale bar, 20 μm. Right: IDR (red) and non-IDR (blue) region of TPM4 according to MobiDB (https://mobidb.bio.unipd.it/) and IUPred (http://iupred.elte.hu). (B) Representative images of purified TPM4-AcGFP protein in storage and phase separation buffer. Scale bar, 10 μm. Zoomed images showing the fusion of TPM4 droplets in vitro. (C) Representative images of TPM4-AcGFP droplets with different protein concentration (0.5, 10, 50 μM) in phase separation buffer. Scale bar, 10 μm. (D) Left: representative images of MDA-MB-231 cells expressing TPM4-mcherry-Cry2 upon blue light exposure and withdrawal. Scale bar, 20 μm. Right: zoomed images showing the formation and fusion of TPM4 condensates upon blue light exposure within 2 min. (E) Representative images of MDA-MB-231 cells expressing TPM4-AcGFP under 100 mM sorbitol treatment and washout. Scale bar, 20 μm. Zoomed images showing the fusion events of TPM4 condensates. (F) Representative time-lapse images of MDA-MB-231 cells expressing TPM4-AcGFP treated with 12.5 mM, 25 mM, 50 mM sorbitol within 2 min. Scale bar, 20 μm. (G) Representative time-lapse images of TPM4-AcGFP and LifeAct-mScarlet in MDA-MB-231 cells treated with 100mM sorbitol within 20 s. Scale bar, 2 μm. (H) Schematic illustration of CRISPR-Cas9-mediated endogenous tagging of TPM4 gene with GFP in C-terminus. (I) Representative images of endogenous TPM4-AcGFP in MDA-MB-231 cell before (isosmotic) and after (hyperosmotic) 100 mM sorbitol treatment for 3 min. Scale bar, 20 μm. (J) Representative images of TPM4 ki in MDA-MB-231 cell before and after 2.5 μM Latrunculin B treatment for 1 h. Scale bar, 20 μm.

We then carried out in vitro phase separation assay using affinity-purified TPM4 protein and observed concentration-dependent droplet formation (Figure 2B and 2C). These droplets underwent rapid fusion when encountering each other (Figure 2B), suggesting a liquid state, which is typical of condensates formed via phase separation. Moreover, we employed an optogenetic approach based on the “optoDroplet” system[37] and found reversible formation of TPM4-Cry2 droplets upon blue light exposure in multiple cell lines (Figure 2D and S1A). These data support the notion that TPM4 can undergo phase separation.

Recent work has shown that hyperosmotic stress could induce phase separation[17]. Following the results that hyperosmotic stress promoted actin remodeling, we asked whether phase separation induced by hyperosmotic stress is involved in this process. Indeed, we observed rapid formation of TPM4 condensates under hyperosmotic stress (100 mM sorbitol; 400 mOsm) (Figure. 2E). These TPM4 condensates are dynamic and reversible upon washing out of the hyperosmotic medium. TPM4 puncta were visible with sorbitol stresses as low as 25 mM (325 mOsm), and the number of puncta per cell increased with higher osmolarity (Figure 2F). Other hyperosmotic stressors exerted similar effects on TPM4 puncta formation (Figure S1B). Co-transfection of TPM4-AcGFP and LifeAct-mScarlet revealed that TPM4 condensates adhered to and wrapped around actin filaments under hyperosmotic stress (Figure 2G and movie 1). Fusion events of TPM4 condensates occurred along actin filaments (Figure S1C and movie 2).

To determine whether these findings are relevant at endogenous levels of TPM4, we generated a TPM4 knockin cell line in which AcGFP was inserted into the C-terminus (referred to as TPM4 ki hereafter, Figure 2H and S1D). When subjected to hyperosmotic medium, the TPM4 ki cells exhibited dramatically enhanced TPM4 localization on actin filaments (Figure 2I). Besides, we noticed that the signal of TPM4 ki and endogenous antibody on actin filaments is discontinuous and condensate-like (Figure 2I and S1E). Interestingly, when we depolymerized the actin with Latrunculin B, which would also release actin-binding TPM4, we observed more pronounced TPM4 condensates in the cytoplasm (Figure 2J).

These results support that TPM4 can undergo phase separation, forming condensates adhering to actin filaments under hyperosmotic stress.

### TPM4 phase separation can recruit glycolytic enzymes

Next, we set out to investigate the role of TPM4 phase separation and its potential relationship with glycolysis. Glycolysis involves at least ten steps, three of which require rate-limiting enzymes PFKM, HK2 (Hexokinase) and PKM2 (Figure 3A). Immunofluorescence staining revealed the presence of PFKM, HK2 and PKM2 within TPM4 condensates upon hyperosmotic stress (Figure 3B). Apart from these, we also identified endogenous signals of other glycolytic enzymes within the TPM4 condensates, including GPI (Phosphoglucose isomerase), ALDOA, TPI (Triosephosphate isomerase), GAPDH (Glyceraldehyde-3-phosphate dehydrogenase), PGK1 (Phosphoglycerokinase), PGAM1 (Phosphoglycerate mutase), and ENO1 (Enolase) (Figure S2A).

**Figure 3.**
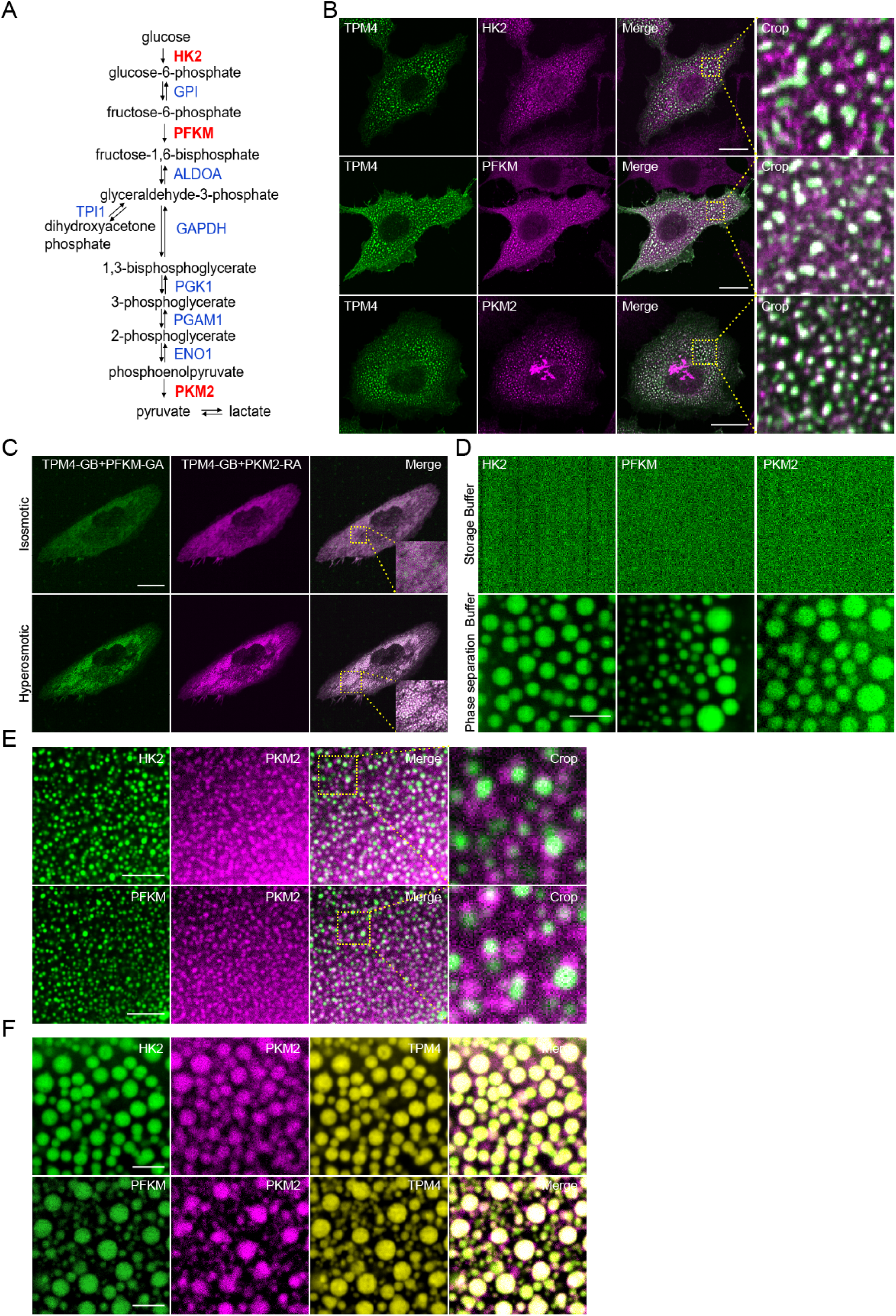
TPM4 condensates recruit rate-limiting glycolytic enzymes. (A) Flow chart showing the steps of glycolysis and key enzymes involved. Three rate-limiting enzymes are highlighted with red. (B) Representative immunofluorescence images stained with HK2, PFKM and PKM2 in MDA-MB-231 cells expressing TPM4-AcGFP under hyperosmotic condition (100 mM sorbitol for 3 min). Scale bar, 20 μm. (C) Representative images in MDA-MB-231 cells expressing TPM4-GB, PFKM-GA and PKM2-RA before (isosmotic) and after (hyperosmotic) 100 mM sorbitol treatment for 3 min. Scale bar, 20 μm. (D) Representative images of purified HK2-AcGFP, PFKM-AcGFP and PKM2-BFP in storage and phase separation buffer. Scale bar, 5 μm. (E) Representative images of purified PKM2-BFP separately with HK2-AcGFP (up) or PFKM-AcGFP (down) proteins in phase separation buffer. Scale bar, 5 μm. (F) Representative co-phase images of purified TPM4-RFP and PKM2-BFP with HK2-AcGFP (up) or PFKM-AcGFP (bottom) in phase separation buffer. Scale bar, 5 μm.

We purified the PFKM, HK2 and PKM2 and observed that all of them could form condensates in vitro respectively (Figure 3D). However, when mixed together, PFKM, HK2 and PKM2 formed distinct condensates (Figure 3E). Interestingly, when added purified TPM4, these glycolytic enzymes formed co-localized condensates (Figure 3F). These results suggest TPM4 may act as a scaffold to organize multiple glycolytic enzymes.

FPX (fluorescent protein exchange) could be used to image dynamic and reversible protein-protein interactions. This system involves two monomers-one (GA or RA) contains a chromophore that is quenched in the monomeric state, while the other (GB) acts only to substantially increase the fluorescence when forming heterodimer (GA/GB or RA/GB) with GA or RA[38]. To visualize the recruitment process of glycolytic enzymes by TPM4 within cells, we fused GA, GB and RA tags to PFKM, TPM4 and PKM2 respectively. Green fluorescence was detected due to the interaction between PFKM-GA and TPM4-GB (Figure 3C upper left) and red fluorescence was detected due to the interaction between TPM4-GB and PKM2-RA (Figure 3C upper middle). Notably, hyperosmotic stress induced the formation of both green and red condensates and they colocalized (Figure 3C bottom and right), supporting that TPM4 can recruit two types of glycolytic enzymes simultaneously.

In conclusion, these findings imply that TPM4 can enrich glycolytic enzymes through phase separation.

### TPM4 phase separation is associated with elevated glycolysis

Having observed that hyperosmotic stress enhanced TPM4 phase separation and recruited glycolytic enzymes, we then asked whether these TPM4 condensates are functionally associated with glycolysis. First, to monitor the production of the glycolysis intermediate NADH produced in the process of Glyceraldehyde 3-phosphate (G3P) converting into 1,3-Bisphosphoglyceric acid (1,3-BPG), we employed the Peredox probe, which contains a bacterial NADH-binding protein Rex and undergoes conformational change upon NADH binding. This shift from an open conformation to a closed one results in the fluorescence change of the linked fluorescent protein[39]. Upon switching into hyperosmotic medium, increased fluorescent intensity was readily observed and resided in a condensed fashion. Notably, the enhanced NADH signal colocalized with TPM4 intensity, as revealed by line scanning (Figure 4A and 4B).

**Figure 4.**
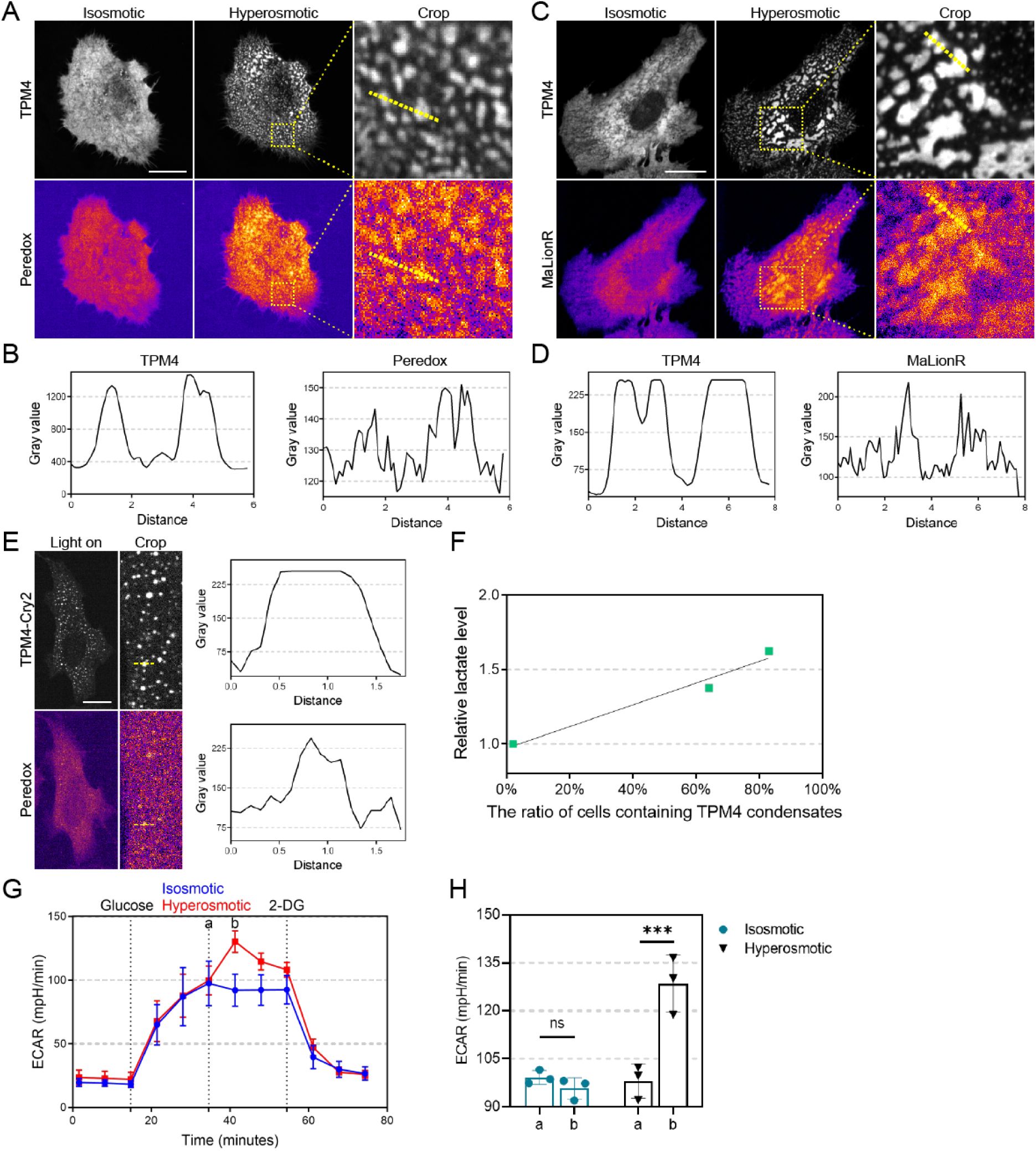
TPM4 phase separation is associated with elevated glycolysis. (A) Representative images of TPM4 and NADH signals by Peredox probe in MDA-MB-231 cells expressing TPM4-AcGFP before (isosmotic) and after (hyperosmotic) 100 mM sorbitol treatment for 3 min. Scale bar, 20 μm. (B) Line scan plot showing the intensity distribution of TPM4 and NADH signals along the yellow line shown in (A) under hyperosmotic condition. (C) Representative images of ATPsignals by MaLionR probe in MDA-MB-231 cells expressing TPM4-AcGFP before (isosmotic) and after (hyperosmotic) 100 mM sorbitol treatment for 3 min. Scale bar, 20 μm. (D) Line scan plot showing the intensity distribution of TPM4 and ATP signals along the yellow line shown in (B) under hyperosmotic condition. (E) Left: representative images of TPM4 and NADH signals in MDA-MB-231 cells expressing TPM4-mcherry-Cry2 upon blue light exposure. Dashed boxes are zoomed in on the right. Scale bar, 20 μm. Right: Line scan plot showing the intensity distribution of TPM4 and NADH signals along the yellow line. (F) Quantification of relative cellular lactate level with the ratio of MDA-MB-231 cells containing TPM4 condensates under hyperosmotic condition. (G) Quantitative curves of ECAR in MDA-MB-231 cells treated with Seahorse basic medium (isosmotic) or 100 mM sorbitol (hyperosmotic) (n = 3 independent experiments, error bar: mean with SEM). Sorbitol was added at time point a, and ECAR reaches its highest level at time point b. (H) Quantification of ECAR in isosmotic and hyperosmotic group at time point a and b (N = 3 independent experiments, error bar: mean with SEM, isosmotic *p*_ab_ = 0.8752, hyperosmotic \*\*\**p_ab_* = 0.0007 by two-way ANOVA).

Second, we sought to monitor ATP level as an indicator of net energy production in cells under hyperosmotic stress. The MaLionR probe, which inserts the ATP-binding ε subunit of the bacterial FoF1-ATP synthase into the red fluorescent protein mApple, changes fluorescence when triggered by ATP binding[40]. Using this probe, we observed that TPM4 condensates induced by hyperosmotic stress exhibited significantly higher levels of ATP compared to surrounding regions (Figure 4C and 4D). These data support TPM4 condensates induced by hyperosmotic stress can serve as hotspots for glycolysis and energy production.

Third, we asked whether opto-genetically induced TPM4 phase separation would be competent to concentrate glycolysis. We shed blue light to TPM4-Cry2 expressing cells and observed that TPM4 condensates showed significantly higher NADH levels compared to surrounding regions (Figure 4E).

Fourth, in support of the notion that hyperosmotic stress triggers functional TPM4-glycolytic enzyme condensation, we detected the overall glycolytic level in cells by two means, one of which was the L-Lactate assay kit. We observed a gradual rise in lactate production by cells associated with a higher ratio of cells with TPM4 condensates (Figure 4F). The other was the Seahorse assay. When we switched normal media to hyperosmotic medium while monitoring bulk glycolysis using the Seahorse, we found that hyperosmotic stress rapidly increased the cellular ECAR (extracellular acidification rate). Subsequently, the use of 2-deoxyglucose (2-DG), a competitive inhibitor of the glycolytic rate-limiting enzyme HK2, significantly reduced ECAR (Figure 4G and 4H). This suggests that the increase in ECAR induced by hyperosmotic stress originates from the glycolytic pathway

Overall, these results support that TPM4 phase separation is functionally associated with elevated glycolysis.

### TPM4 mediates actin reorganization coupled with glycolysis regulation

We observed that TPM4 phase separation facilitated the recruitment of glycolytic enzymes onto the actin filaments, promoting glycolysis. An intriguing question arises: what are the metabolic consequences of TPM4 absence? To answer this, we generated a TPM4 knockout cell line by CRISPR/Cas9 (Figure S3A). In normal cells under hyperosmotic stress, there was colocalization of the PFKM and actin filaments (Pearson’s coefficient = 0.486), as evidenced by the alignment of their distribution patterns in line scanning (Figure 5A); however, this correlation was not observed in TPM4 KO cells (Pearson’s coefficient = 0.143) (Figure 5B). Following this, we observed that the elevation levels of lactate in the TPM4 KO group were lower compared to the control group under hyperosmotic stress. (Figure S3B). These observations argue that TPM4 is essential for maintaining glycolytic activity under hyperosmotic stress.

**Figure 5.**
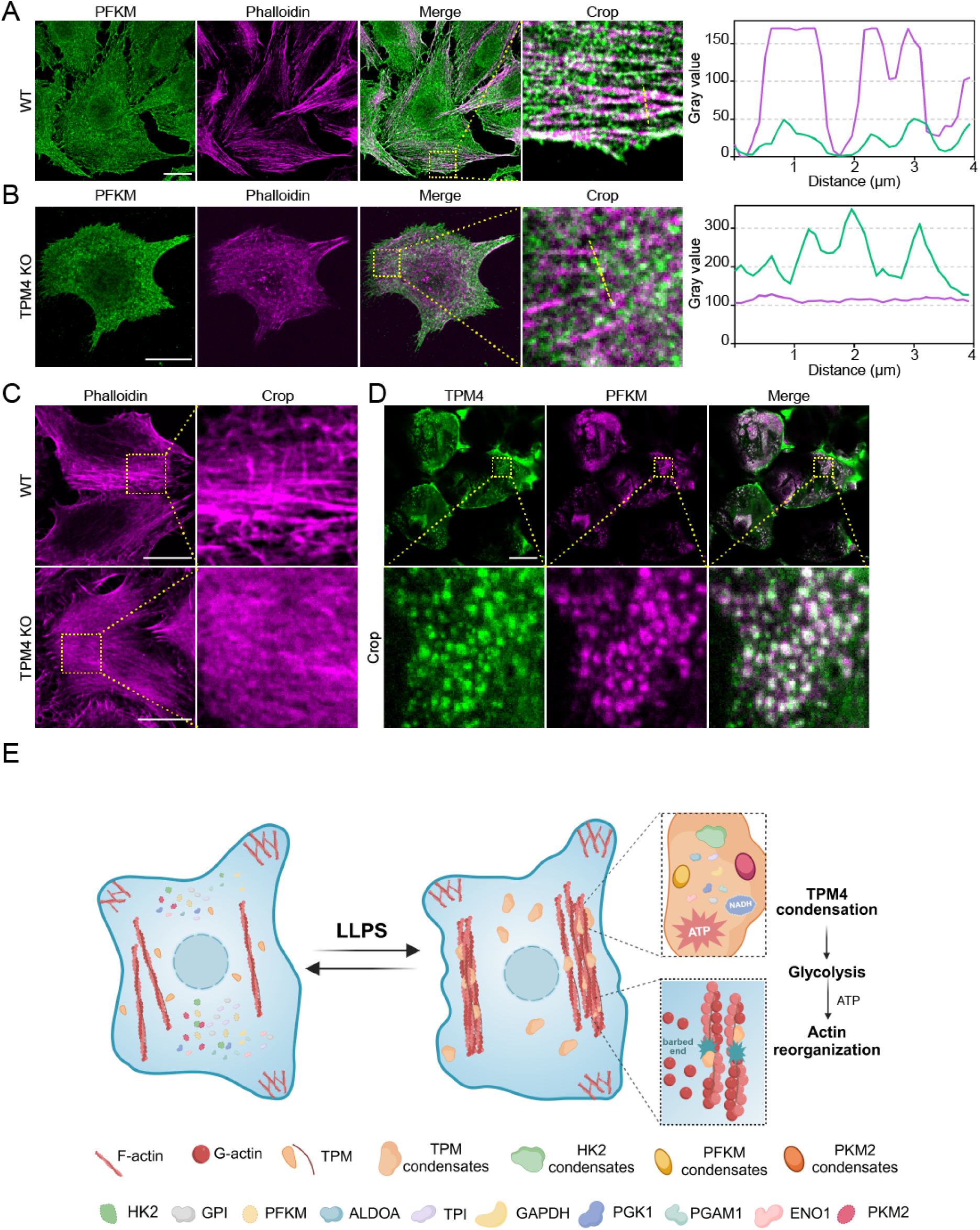
TPM4 mediates actin reorganization coupled with regulation of glycolysis. (A) Left: Representative immunofluorescence images stained with PFKM (green) and phalloidin (magenta) in MDA-MB-231 WT cells treated with 100 mM sorbitol for 3 min. Scale bar, 20 μm. Right: line scan plot showing the intensity distribution of PFKM and actin filaments along the yellow line. (B) Representative immunofluorescence images stained with PFKM (green) and phalloidin (magenta) in MDA-MB-231 TPM4 KO cells treated with 100 mM sorbitol for 3 min. Scale bar, 20 μm. Right: line scan plot showing the intensity distribution of PFKM and actin filaments along the yellow line. (C) Representative immunofluorescence images stained with phalloidin (magenta) in WT and TPM4 KO cells treated with 100 mM sorbitol for 1 h. Scale bar, 20 μm. (D) Representative immunofluorescence images stained with TPM4 (green) and PFKM (red) in mouse kidney tissue sections. Scale bar, 20 μm. (E) A model for TPM4 phase separation mediated reorganization of multiple glycolytic enzymes. Upon hyperosmotic stress, TPM4 undergoes phase separation, recruiting 10 essential glycolytic enzymes to upregulate glycolysis and support energy with actin reorganization.

In TPM4 KO cells, no significant changes were observed in actin filaments under isosmotic conditions (Figure S3C). However, under hyperosmotic stress, less and thin actin filaments appeared in TPM4 KO cells compared with normal cells (Figure 5C). This finding support that the TPM4 phase separation and its recruitment of glycolytic enzymes onto actin filaments are important for actin reorganization under hyperosmotic stress.

### TPM4-PFKM condensates can be observed in physiological conditions

Many organ tissues within the body, including but not limited to renal, hepatic, and intestinal tissues, establish and maintain osmotic homeostasis. The renal tissue sustain osmotic stress as high as 1200 mOsm[41]. This raises an intriguing question: whether inherently hyperosmotic environment in renal tissues triggers TPM4 condensation? Indeed, by immunofluorescence staining we detected TPM4 condensates in mouse kidney tissues (Figure 5D). Moreover, these condensates also displayed strong colocalization with PFKM (Figure 5D).

In summary, our data suggest a model that actin reorganization is coupled to glycolysis through phase separation under hyperosmotic stress (Figure 5E): the actin-binding protein TPM4 can undergo phase separation, forming condensates that adhere to and wrap around actin filaments. TPM4 condensates are able to recruit multiple essential glycolytic enzymes including HK2, PFKM and PKM2, promoting glycolysis and subsequently facilitating actin reorganization.

## Discussion

Dynamic rearrangements of the actin cytoskeleton are critical for various cellular processes, including morphological change, cell motility, mechanical integrity and transcription regulation[42]. Studies on unstimulated platelets revealed that approximately 50% of total ATP consumption is dedicated to maintaining actin dynamics [1]. Similarly, in neurons, when actin turnover is inhibited, either by blocking actin disassembly or assembly, ATP depletion is reduced by 50%[2]. How does actin cytoskeleton acquire such a significant amount of ATP in time scales of rapid reorganization? Early indications that glycolysis may provide the necessary fuel for the actin cytoskeleton have emerged from studies reporting direct interactions between glycolytic enzymes and actin. For instance, phosphofructokinase-1 (PFK-1) binds to actin through electrostatic interactions in vitro[3]. Aldolase has been shown to preferentially bind to F-actin rather than G-actin in vitro[4]. Furthermore, GAPDH also directly interacts with F-actin in vitro[5]. Our observations revealed that upon exposure to hyperosmotic stress, the actin-binding protein TPM4 can undergo phase separation and recruit ten essential glycolytic enzymes. TPM4 condensates had high levels of glycolytic intermediates and ATP, suggesting that TPM4-glycolytic enzyme condensates on the actin filaments may efficiently supply energy for the dynamic actin rearrangement under hyperosmotic stress. It would be interesting to explore the energy sources and potential mechanisms of actin cytoskeleton rearrangement under other stress conditions.

Subunit assembly and disassembly constantly remodel actin filaments. In our study, TPM4 undergoes phase separation, forming condensates that adhere to actin filaments. When abundant actin filaments are present in the cell, TPM4 tends to undergo phase separation on the filaments. However, when actin filaments are depolymerized, TPM4 loses its attachment sites on the filaments, resulting in dispersed TPM4 condensates throughout the cytoplasm. Hyperosmotic stress induces the perturbation of actin filaments, leading to the generation of more free barbed ends in the cell.

Normally, the ten core glycolysis enzymes are diffusely localized throughout the cytoplasm[43]. Different components could be brought together through LLPS to react efficiently[18, 19]. Thus, LLPS may act as a platform for glycolysis. We found that TPM4 phase separation can recruit multiple glycolytic enzymes, mediating the compartmentalization of glycolytic enzymes. This compartmentalization may enhance the efficiency of glycolysis, meeting the energy demands of actin dynamics. When actin assembly requires a quick energy supply, actin-binding protein TPM4 undergoes phase separation and recruits glycolytic enzymes to provide energy. Similarly, when other cellular structure organization requires energy, could corresponding core proteins recruit glycolytic enzymes to supply energy? Alternatively, the ubiquitous actin cytoskeleton and its associated TPM4-glycolytic enzyme condensates may not only provide energy to actin filaments but also satisfy local cellular energy demands.

Some glycolysis-related condensates have been reported in the literature. For instance, “glucosome” is an assembly of glycolytic enzymes including PFK, FBPase, PEPCK, and PKM, organized to regulate glucose flux in human cancer cells[44]. Additionally, under hypoxic stress, yeast and C. elegans neurons form “G bodies” or “glycolytic granules” containing enzymes like PFK, potentially enhancing glycolytic activity and ATP production[23–25]. In our study, we discovered that TPM4 condensates formed through phase separation localize on the actin filaments and enrich essential glycolytic enzymes including HK2, PFKM and PKM2. TPM4-mediated glycolytic compartmentalization thus promotes glycolysis and facilitate actin reorganization, providing a novel model that links actin cytoskeleton and glycolysis. In general, these different glycolysis-related condensates may represent strategies for cells to adapt to various environmental stresses.

Limitation of this study: Our in vivo studies on hyperosmotic stress-induced TPM4 phase separation are relatively limited, and the presence of TPM4 condensates has mainly been demonstrated through simply staining of mouse kidney tissues. However, there is still a lack of substantial evidence regarding whether TPM4 in renal cortex cells and medullary cells exhibits different organizational modes and physiological functions under different osmotic environments. Further experiments are needed to observe the responses of TPM4 in different cell types within the kidney to varying osmotic stresses and determine their respective physiological functions. In our study, we focused on the spatial recruitment of various glycolysis-related enzymes by TPM4 after hyperosmotic stress but lacked the detection of enzyme activity in condensates. Exploring the activities of different enzyme components in condensates formed after phase separation is crucial for functional studies and represents a challenging and important research direction in the field of phase separation.

## Methods

### Cell lines and cell culture

Human embryonic kidney 293T (HEK 293T) cells were kept by our laboratory. MDA-MB-231 cells were generously provided by Yujie Sun laboratory (Peking University, China). Cells were cultured in Dulbecco’s modified Eagle medium (DMEM; Corning, 10-013-CRVC) supplemented with 10% fetal bovine serum (FBS; PAN-Biotech, P30-3302), 100 U/mL penicillin and 100 μg/mL streptomycin at 37°C with 5% CO_2_. For cell passage, cells were washed with PBS (Macgene, CC010) and digested with trypsin (Macgene, CC012).

### Plasmids and transient infection

Passage the cells 12 h prior to plasmid transient transfection to ensure that the cell density for transfection falls within the 50% to 60% range. About 1 h before transfection, replace the medium with fresh cell culture medium. Prepare the plasmid transient transfection system as per the instructions. After mixing, allow it to stand for 20-30 min before evenly adding it to the culture dish. Around 12 h later, replace with fresh medium. The transfected plasmids in the cells exhibit high expression levels within 36-72 h, enabling live-cell imaging, immunofluorescence staining, and protein immunoblotting experiments.

### CRSPR/Cas9-mediated TPM4 gene knockout

For gene knockout, the following sgRNAs in LentiCRISPR-V2 (#52961, Addgene) were used in MDA-MB-231 cells. After puromycin selection, the single cell clones were cultured and verified by sequencing and western blotting.

For TPM4 knockout:

sgRNA-1: 5’-cgagctggataaatattccg-3’

sgRNA-2: 5’-gcttctcctgcgcgtccttc-3’

### Generation of TPM4-AcGFP knock-in cells

The following sgRNAs were cloned into lentiCRISPR-V2 vector. The donor templates contained left homologous arm (800 bp before the stop codon), AcGFP sequence and right homologous arm (800 bp after the stop codon). The homologous arms were amplified from the MDA-MB-231 cell extracted genome DNA. To generate the knock-in cells, sgRNA and relative donor template plasmid were co-transfected to the cells. After 24h, puromycin was used to eliminate the puromycin-sensitive cells. The fluorescence positive cells were acquired by Fluorescence Activated Cell Sorting (FACS) and the single cell clones were cultured and verified by western blotting.

sgRNA for TPM4 C-terminal knock-in in MDA-MB-231 cells: 5’-tggtgaggattaagagtatg-3’.

### Prediction of disordered tendency

The intrinsically disordered tendency of TPM4 was predicted by MobiDB (https://mobidb.bio.unipd.it/) and IUPred (http://iupred.elte.hu).

### Western blotting

For western blotting, cells were washed with PBS and lysed in an appropriate volume of RIPA buffer (50 mM Tris, pH 8.0, 150 mM NaCl, 1% Triton X-100, 0.5% Na-deoxycholate, 0.1% SDS,1 mM EDTA and protease inhibitor cocktail) for 10 min on ice. Lysates were centrifuged at 13,572 g for 10 min and the supernatants were collected. Then, 5 × SDS loading buffer was added to the supernatants and boiled for 10 min. Protein samples were run on 10% SDS–PAGE acrylamide gels and transferred onto nitrocellulose (NC) membranes by wet electrophoretic transfer, followed by primary and secondary antibody incubation at 4°C overnight or at room temperature for 2 h. Use a gel imaging system for result collection. Prepare a working solution by mixing luminescent liquid A and B at a 1:1 ratio, then apply it evenly onto the strip. After a 2 min incubation, directly perform exposure imaging on the gel imaging instrument, adjusting the exposure time based on the depth of the strip. The band intensity was quantified by ‘gel analysis’ plugin of Fiji software (https://fiji.sc).

### Proximity labeling with TurboID

Seed MDA-MB-231 cells into a 6 cm cell culture dish, then perform transient transfection of the plasmids Plvx-TPM4-Turbo, Plvx-PDLIM1-Turbo, Plvx-ACTN1-Turbo, Plvx-ArgBP2-Turbo, and Plvx-Turbo. After 24 h, passage the cells into a 10 cm cell culture dish respectively. Then use biotin for labeling at a working concentration of 50 μmol/L. Discard the original culture medium, add prepared biotin-containing medium, and incubate for 10 min. After that, remove the culture medium, wash 2-3 times with cold PBS to remove residual biotin, Protein was extracted using lysis buffer (RIPA, 50 mM Tris-HCl, pH 8.0, 150 mM NaCl, 1% Triton X-100, 0.5% Na-deoxycholate, 0.1% SDS, 1 mM EDTA and protease inhibitor cocktail). Thoroughly mix the streptavidin magnetic bead suspension, pipette 50 μL of the magnetic bead suspension, and then use a magnetic stand to collect and wash the beads with lysis buffer 3 times. Take 1 mg of protein and thoroughly mix it with the magnetic bead suspension, followed by incubation overnight at 4°C on a shaking bed. The following day, wash the overnight incubated magnetic beads twice with lysis buffer, followed by washing with 1 mol/L KCl solution, 0.1 mol/L Na_2_CO_3_ solution, and 2 mol/L urea (10 mM Tris-HCl, pH 8.0), then perform two additional washes with RIPA lysis buffer, ensuring each step is completed promptly. Prepare mass spectrometry buffer (0.1% SDS in 55 mM Tris-HCl, pH 8.0) and add 6.66 mM DTT and 0.66 mM of biotin. Add 120 μL of the prepared buffer to the washed magnetic beads, mix well, and boil in a 98°C metal bath for 10 min. Use 20 μL of the protein for protein immunoblotting and store the remaining 100 μL of protein at −80°C for mass spectrometry identification.

### Immunofluorescence and imaging analysis

Cells were plated on acid-washed coverslips coated with 10 mg/mL fibronectin overnight. Cells were then fixed with 4% paraformaldehyde (PFA) at room temperature for 20 min, permeabilized in 0.5% Triton X-100 in PBS for 5 min, washed with PBS once for 5 min and blocked with 10% bovine serum albumin (BSA) for 1 h. Then the primary antibody was diluted 1:200 in PBS and incubated for 1 h at room temperature. After washing with PBS 3 times, the coverslips were incubated with Alexa Fluor 488 or 555-conjugated secondary antibody (1:200) for 1 h at room temperature. After another wash with PBS 3 times, the coverslips were mounted with ProLong^TM^ Glass Antifade Mountant with NucBlue^TM^ Stain (P36981, Invitrogen). After mounting medium was solidified, images were captured by Andor Dragonfly confocal imaging system.

### Kidney dissection and immunofluorescence

Animal studies were conducted according to guidelines approved by the Biomedical Ethics Committee of Peking University (approval no. LA2020398). Kidney tissues from two wild-type C57BL/6-8 weeks mice were immersion-fixed with 4% paraformaldehyde (PFA) in PBS for 48 h at 4°C. For the immunofluorescence microscopy staining, embedding the kidney tissue using the frozen section embedding medium OCT and sectioned into 10μm thick slices using a vibratome (Leica). Expose the sections to room temperature for 2-3 h to ensure the tissue fully adheres to the glass slide and prevent detachment. Then, fix with 4% paraformaldehyde for 30 min at room temperature. Wash the sections 2-3 times with PBS, followed by permeabilization with 0.5% Triton-X 100 for 30 min at room temperature. Transfer the permeabilized sections to a humidified chamber, use an immunohistochemistry pen to draw a circle slightly larger than the tissue around the tissue, then block with 10% BSA containing 0.5% Triton-X 100 at room temperature for 1 h. The sections were incubated sequentially with a primary antibody against TPM4 (1:100; Proteintech) and PFKM (1:100, Abclonal) in buffer (0.5% Triton-X 100 and 2% BSA). for 2 h. After washing with PBS 3 times, the coverslips were incubated with Alexa Fluor 488 or 555-conjugated secondary antibody (1:200) and phalloidin (1:400) for 1 h at room temperature. After another wash with PBS 3 times, the coverslips were mounted with ProLong^TM^ Glass Antifade Mountant with NucBlue^TM^ Stain (P36981, Invitrogen). After mounting medium was solidified, images were captured by Andor Dragonfly confocal imaging system.

### Live cell imaging of Opto-droplet system

For long-term live cell imaging, cells were plated on 10 mg/mL fibronectin coated glass-bottom cell culture dishes and maintained in CO_2_-independent DMEM (Gibco, 18045-088) supplemented with 10% FBS, 100 U/mL penicillin and 100 mg/mL streptomycin at 37°C throughout the imaging process. For Opto-droplet phase separation imaging, cells were covered with tin foil to block light and images were acquired at indicated intervals using a 63 × 1.4 NA objective lens on Andor Dragonfly confocal imaging system.

### Live cell NAhDH and ATP determination

For live cell imaging of NADH and ATP determination, cells were transfected with the NADH biosensor “Peredox-mCitrine (λ_ex_/λ_em_ = 400/510 nm)” and the ATP biosensor “MaLionR” (λ_ex_/λ_em_ = 565/585 nm). These cells were plated on 10 mg/mL fibronectin-coated glass-bottom cell culture dishes and maintained in CO_2_-independent DMEM (Gibco, 18045-088) supplemented with 10% FBS, 100 U/mL penicillin, and 100 mg/mL streptomycin at 37°C throughout the imaging process.

### Proximity ligation assay (PLA)

PLA was performed with Duolink kits from Sigma-Aldrich. Cells were then fixed with 4% paraformaldehyde (PFA) at room temperature for 20 min. After 3 times PBS washing, the cells were permeabilized with 0.5% Triton X-100 in PBS for 5 min. Commercial blocking solution was added to the samples and incubated for 1 h at room temperature.

After blocking, the cells were incubated with the diluted antibodies for 1 h at room temperature followed by 3 times PBS washing. The PLUS and MINUS PLA probes were mixed and diluted (1:5) in antibody diluent and incubated with samples for 30 min at 37°C. Then, the samples were washed in 1 × Wash Buffer A for 5 min twice. The ligase was diluted (1:40) in ligation buffer (1:5 diluted in H_2_O) and incubated with samples for 30 min at 37°C, followed by washing with 1 × Wash Buffer A for 2 min twice. The polymerase was diluted (1:80) in amplification stock (1:5 diluted in H_2_O) and incubated with samples for 100 min at 37°C. The samples were then washed in 1 × Wash Buffer B for 10 min twice, followed by another washing in 0.01 × Wash Buffer B for 1 min. Finally, the samples were mounted with Prolong Diamond Antifade with DAPI for 30 min at room temperature. For positive control of PLA experiments, anti-tyrosinated α-tubulin and anti-α-tubulin antibodies were used to identify the positive signals. For negative control of PLA experiments, anti-PFKM or anti-GFP antibodies were used alone.

### Constructs, protein expression and purification

For recombinant protein expression in E. coli BL21 (DE3) cells, DNA fragments were cloned into the pET28a (+) vector. Full length TPM4-AcGFP, TPM4-RFP, PFKM-AcGFP, HK2-AcGFP and PKM2-BFP were cloned into pET28a (+) vector with N-terminal 6 × His tag. The related plasmids were transformed into E. coli BL21 (DE3) cells and protein expression was induced with 100 μM IPTG at 18 °C. To determine whether the protein was expressed in the inclusion body, the cells expressed proteins were lysed in PBS supplied with 1% Triton-X100 and 1 mM phenylmethanesulfonylfluoride (PMSF) using ultrasonic cell crusher and centrifuged at 13,572 g for 10 min. The supernatant and pellet were collected for western blotting. Then the cells expressed proteins were lysed in 1 mM Tris (2-carboxyethyl) phosphine (TCEP) and 1 mM phenylmethanesulfonylfluoride (PMSF) in PBS using ultrasonic cell crusher and centrifugation at 48,380 g for 30 min. The supernatant was applied to a Ni-IDA beads (Smart Lifesciences) and washed with buffer containing 1mM TCEP with 20mM imidazole. After that, proteins were eluted with elution buffer containing 1mM TCEP and 300 mM imidazole in PBS. Proteins were concentrated by centrifugation at 3000 g at 4 °C until reaching the volume of 500 μL using 10 kDa concentrator (UFC9010, Sigma-Aldrich), and then loaded onto a Superdex200 Increase 10/300 (GE Healthcare) equilibrated with gel filtration buffer containing 20 mM Tris, pH 8.0, 500 mM NaCl, and 1 mM TCEP. Peaks containing proteins were collected and evaluated with Coomassie blue staining of SDS-PAGE gels.

### In vitro phase separation assay

The purified proteins are typically stored in a storage buffer (high-salt concentration, 500 mM NaCl, 20 mM Tris-HCl, pH 8.0), which inhibits protein phase separation. When conducting in vitro phase separation experiments, it is necessary to dilute the proteins according to the desired concentration, ensuring that the ultimately diluted proteins can undergo phase separation experiments in a normal salt concentration phase separation buffer (150 mM NaCl, 20 mM Tris-HCl, pH 8.0). Depending on the characteristics of the target protein, metal ions such as Mg^2+^ and Zn^2+^, as well as dextran, can be included. The prepared phase separation system is introduced into a self-made phase-separation microfluidic channel and after standing, image acquisition is performed using a 63 ×1.4 NA objective lens on Andor Dragonfly confocal imaging system.

### Detection of lactate levels

The lactate levels were measured using the Lactate Assay Kit from BioVision (ab65330/K607-100). Lactate can react with the enzymes provided in the kit, producing a noticeable absorbance at 570 nm, which can be used to assess lactate levels.

In brief, following treatment of the cells with 50 mM sorbitol or PBS for 10 min, the cells were digested, and approximately 2 × 10^6^ cells were suspended in DBPS, followed by two washes with PBS. Following the washes, Lactate Assay Buffer from the kit was added to lyse the cells, and the lysis was carried out on ice for 20 min, with gentle tapping of the centrifuge tube to ensure thorough lysis. Following lysis, the lysate was centrifuged at 4°C, 12,000 rpm for 10 min, and the supernatant was transferred to a new centrifuge tube for further use. The reaction system (96-well cell culture plate) was prepared according to the kit instructions, mixed well, kept away from light, and incubated at 37°C for 30 min. Upon completion of the reaction, the absorbance at 570 nm was measured using a microplate reader.

### Seahorse Assay

An XF extracellular flux analyzer (Seahorse Bioscience) was used to determine the effects of the hyperosmotic stimulation on MDA-MB-231 cells, which were seeded at 20,000 cells/well. Measurements of ECAR were performed according to the manufacturer’s instructions.

In brief, The Seahorse XF Detection System needs to be powered on in advance for at least 5 h of preheating. After seeding cells using the XF Cell Culture Microplate, add 80 μL of cell suspension to each well, approximately 2 × 10^4^ cells, to achieve a confluency of around 90% after cell attachment. Allow the cell suspension to settle for 1 h before gently placing it into a 37°C cell culture incubator. Equilibrate the Seahorse sensor cartridge with double-distilled water and then incubate it overnight in the 37°C cell culture incubator. Following this, perform a second equilibration using preheated XF Calibrant Solution, also in the 37°C cell culture incubator for 60-70 min.

In addition to glucose and 2-deoxyglucose used in the Glycolysis Stress Test Kit, this study related to cellular energy metabolism involves assessing the impact of hyperosmotic stimulation induced phase separation on cellular glycolysis. Therefore, Sorbitol solution needs to be added during drug administration, with the control group using the Seahorse assay medium. The drug addition method involves adding a fixed volume of each drug: drug port A contains a glucose solution at a final concentration of 10 mM; drug port B contains a sorbitol solution at a final concentration of 50 mM, or the Seahorse assay medium for the control group; and drug port C contains a 2-deoxyglucose solution at a final concentration of 100 mM. Upon drug addition, an equal volume of the same drug should be added to the experimental and control wells in drug ports A, B, and C. Following the addition of drugs, the system is ready for subsequent detection.

### Statistics and data display

The number of biological and technical replicates and the number of samples are indicated in figure legends, the main text and the STAR methods. The mass spectrometry data is processed using Excel, and subsequently enriched pathway analysis is conducted on the Metascape website (https://metascape.org/gp/index.html#/main/step1). Seahorse’s results are analyzed and processed using the desktop version of WAVE software. Data are mean ± SEM as indicated in the figure legends and supplementary figure legends. Student’s t-test, two-way ANOVA analysis were performed with GraphPad Prism 8.0 and Excel (Microsoft). Data from image analysis was graphed using Prism 8.0.

---

**Figure S1.**
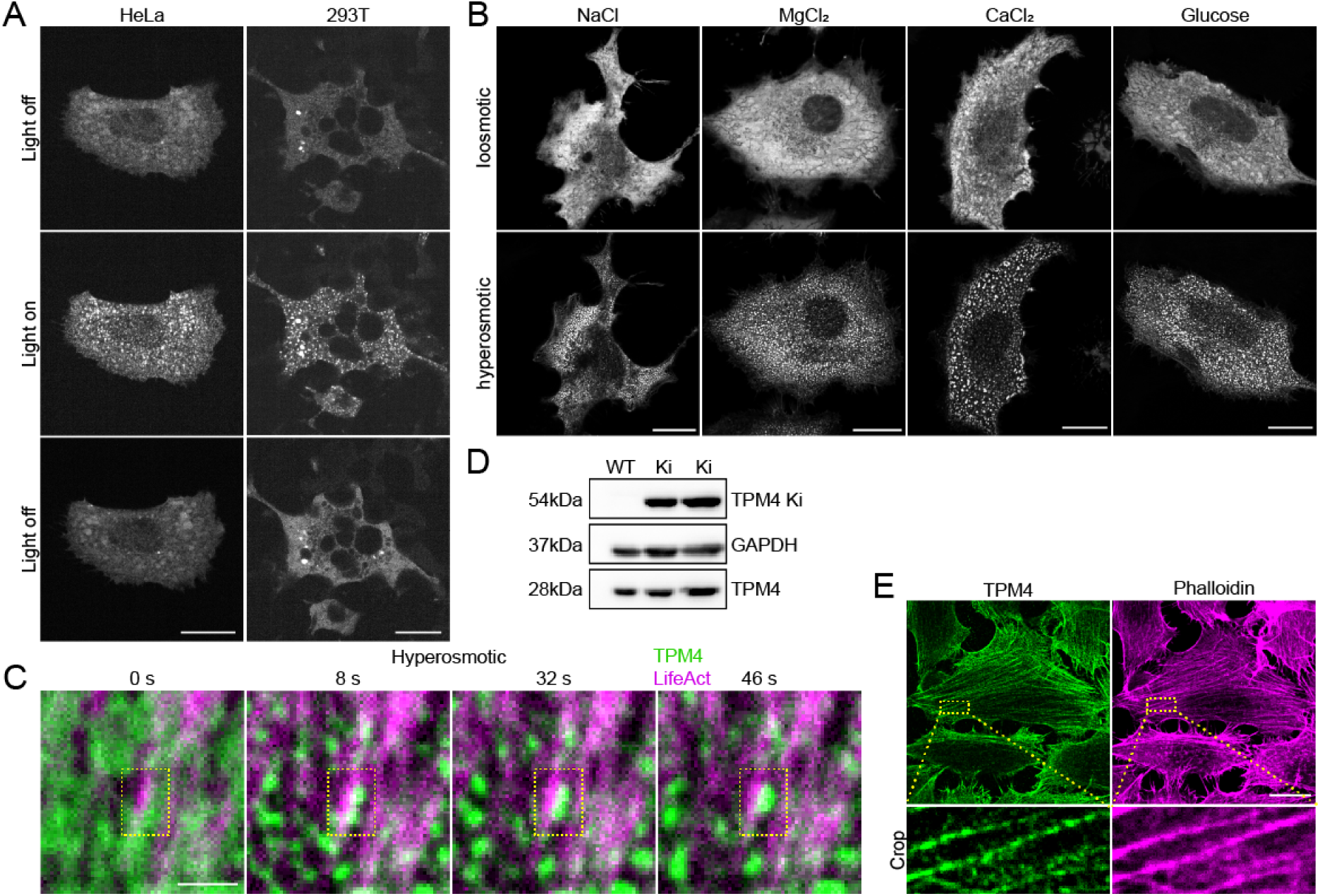
TPM4 condensates formation is dynamic and correlate with actin filaments. (A) Representative images of TPM4-Cry2 in HeLa and HEK 293T cells upon blue light exposure and withdrawal. Scale bar, 20 μm. (B) Representative images of TPM4-AcGFP before and after treatment with 100 mM NaCl, 100mM MgCl_2_, 100mM CaCl_2_ or 100 mM glucose for 3 min in MDA-MB-231 cells. Scale bar, 20 μm. (C) Representative time-lapse images of TPM4 condensates formation along actin filaments (labelled by LifeAct) in 100mM sorbitol treated MDA-MB-231 cells. Scale bar, 2 μm. (D) Western blot showing the successful knockin of TPM4-AcGFP in MDA-MB-231 cells. GAPDH is used as loading control. (E) Representative images of endogenous TPM4 antibody and actin filaments (stained by phalloidin). Scale bar, 20 μm.

**Figure S2.**
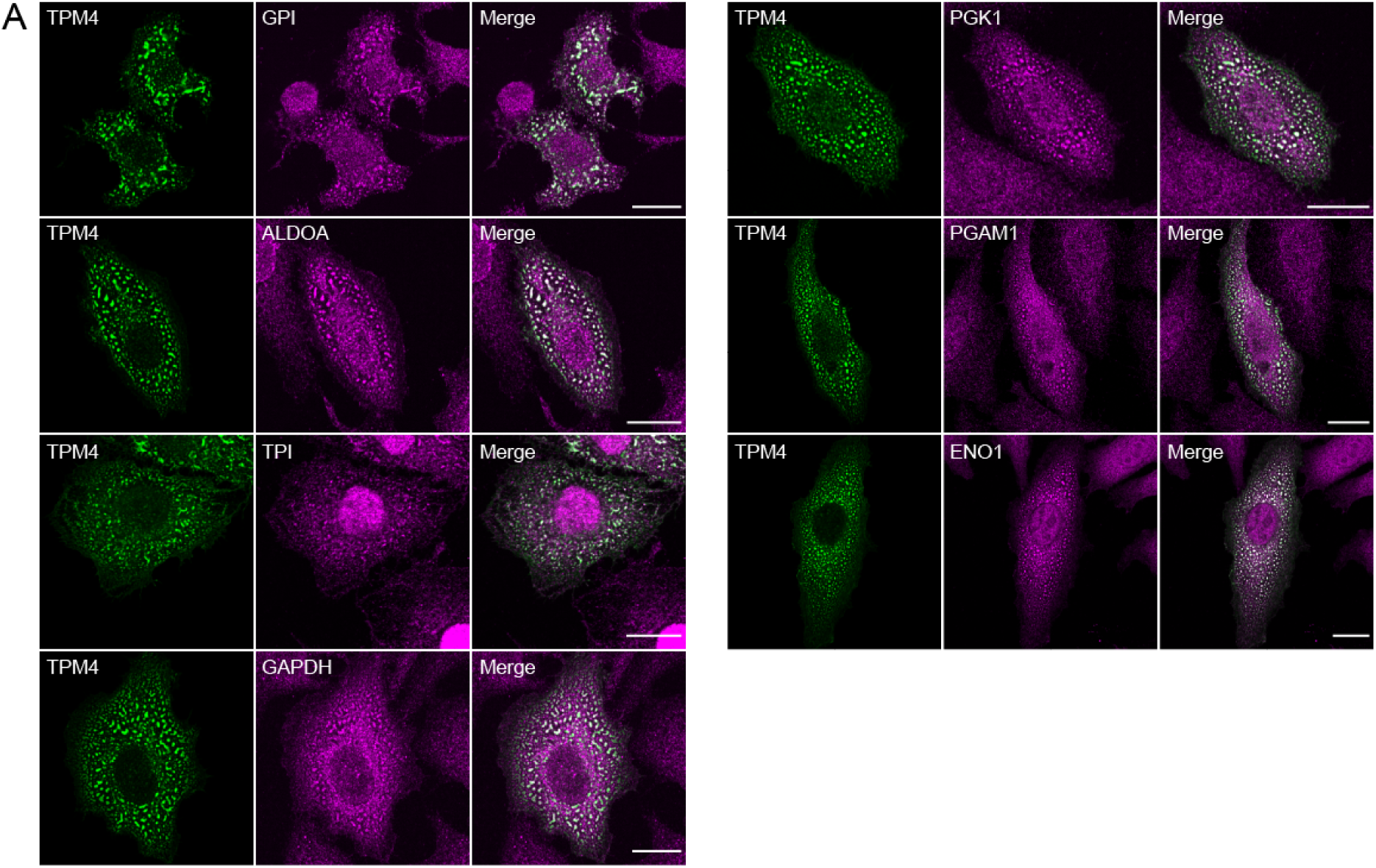
TPM4 condensates contain multiple glycolytic enzymes. (A) Representative immunofluorescence images stained with GPI, ALDOA, TPI, GAPDH, PGK1, PGAM1 and ENO1 in MDA-MB-231 cells expressing TPM4-AcGFP under hyperosmotic condition (100 mM sorbitol for 3 min). Scale bar, 20 μm.

**Figure S3.**
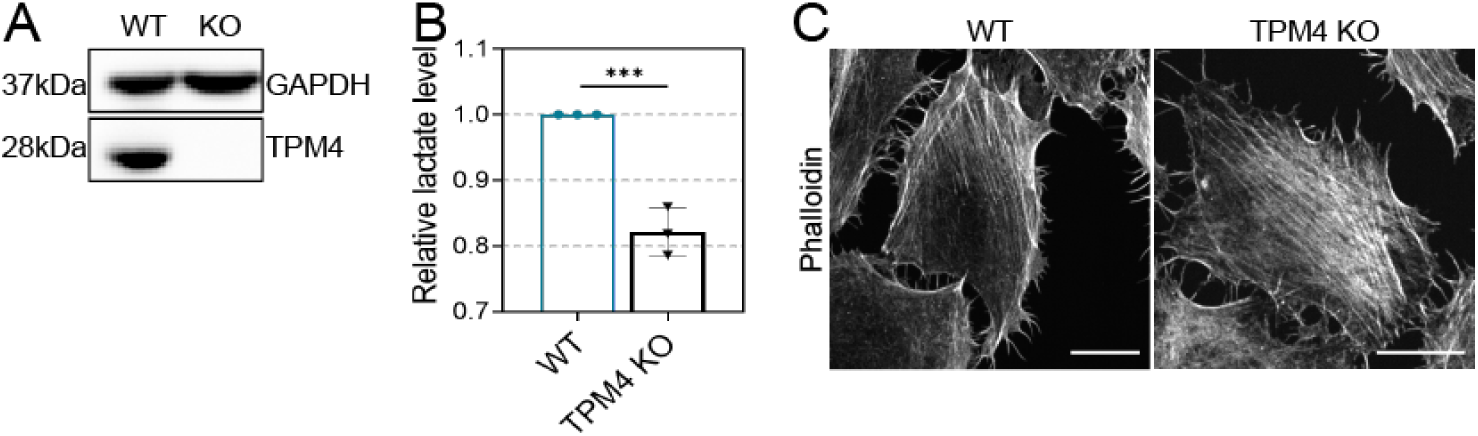
TPM4 KO does not have a significant effect on actin filaments under isosmotic conditions. (A) Western blot showing the successful knockout of TPM4 in MDA-MB-231 cells. GAPDH is used as loading control. (B) Quantitation of relative lactate level in WT and TPM4 KO cells with 100mM sorbitol treatment for 10 min (n = 3 independent experiments, error bar: mean with SEM, \*\**p* = 0.0011 by unpaired *t* test). (C) Representative immunofluorescence images stained with phalloidin in WT and TPM4 KO cells. Scale bar, 20 μm.

## Notes

### Competing Interest Statement

The authors have declared no competing interest.

### Summary of Updates

We have revised the title and abstract of the article.

